# Early Transcriptomic Signatures of Reductive Stress Cardiomyopathy Reveal Insufficient HIF-1α Signaling

**DOI:** 10.1101/2025.09.01.672999

**Authors:** Rishitha Chowdary Mavillapalli, Janani Dakshina Moorthy, Yazhini Ravi, Namakkal S. Rajasekaran

## Abstract

Chronic activation of Nrf2-regulated antioxidant pathways can lead to a pathological condition known as reductive stress (RS). While transient Nrf2 activation confers protection against oxidative stress, its sustained stimulation may provoke maladaptive responses in high-energy-demanding organs such as the heart. In this study, we employed a cardiac-specific constitutively active Nrf2 transgenic (CaNrf2-TG) mouse model to characterize early transcriptomic alterations associated with the transition from compensated cardiac function to pathological remodeling. Myocardial RNA sequencing at 12 weeks of age, a critical window preceding overt cardiac dysfunction, revealed widespread dysregulation of antioxidants, immune, metabolic, proteostasis, and stress-response pathways. Notably, genes related to HIF-1 signaling, PI3K-Akt, MAPK, and hypertrophic cardiomyopathy were significantly altered. Functional enrichment analysis highlighted upregulation of detoxification enzymes and stress response regulators, alongside downregulation of ER chaperones, calcium-handling proteins, and MHC-II immune mediators. Furthermore, perturbations in metabolic flexibility and sarcomeric gene expression suggest early disruption of structural and energy-regulating networks. These findings uncover early molecular signatures of Nrf2-driven RS cardiomyopathy and may aid in identifying potential therapeutic targets for mitigating disease progression.

## 1. Introduction

Cardiovascular diseases (CVDs) remain the leading cause of morbidity and mortality worldwide (1). For decades, oxidative stress has been recognized as a central mechanism in the onset and progression of cardiac dysfunction (2). More recently, an opposing yet equally pathological condition, reductive stress (RS), has gained attention for its deleterious impact on cardiac health (3-9). RS arises from an excessive accumulation of reducing equivalents such as reduced glutathione (GSH), NADH, and NADPH, resulting in hyper-reductive intracellular conditions that disrupt redox signaling and cellular homeostasis. Unlike oxidative stress, RS impairs endoplasmic reticulum (ER) protein folding, alters proteostasis, and initiates maladaptive cardiac remodeling.

While transient activation of nuclear factor erythroid 2–related factor 2 (NFE2L2 or Nrf2) confers cardio protection (10), chronic or constitutive Nrf2 signaling has been implicated in RS (11). In cardiac-specific transgenic mice expressing constitutively active Nrf2 (CaNrf2-TG), sustained Nrf2 activity leads to persistent antioxidant gene expression and accumulation of reducing equivalents, disrupting ER proteostasis and initiating maladaptive signaling cascades (11). Previous studies from our group have shown that CaNrf2-TG hearts display progressive cardiac remodeling characterized by upregulation of antioxidant genes, downregulation of ER chaperones (*Hspa5, Calr*), and activation of the ER-associated degradation (ERAD) pathway (4, 5).

Despite growing recognition of RS and its impact on cardiac health, the early molecular events underlying the shift from adaptive to maladaptive Nrf2-driven transcriptional signaling remain poorly defined. In the present study, we employed RNA sequencing and integrative pathway enrichment analysis to define the early transcriptomic landscape of CaNrf2-TG hearts at 12 weeks. Our findings reveal widespread dysregulation of pathways associated with HIF-1 signaling, hypertrophic cardiomyopathy, sarcomere and mitochondrial function, and proteostasis. These data shed light on the transcriptional mechanisms by which sustained Nrf2 activation contributes to RS-induced cardiac dysfunction and highlight potential early targets for therapeutic intervention.

## 2. Experimental Methods (at the end)

**(Brief methods only and detailed methods will be presented as supplemental section)**

## 3. Results

### 3.1 Constitutive Nrf2 activation drives distinct transcriptomic reprogramming before the onset of cardiac remodeling

Figure 1A presents principal component analysis (PCA) of myocardial transcriptomes from wild-type (WT) and CaNrf2-TG mice at 12-weeks of age. The first principal component (PC1) accounts for 96.26% of the total variance, while PC2 explains 1.17%, indicating a clear separation between the two groups. This substantial divergence suggests that constitutive activation of Nrf2 significantly alters the myocardial transcriptome. Figure 1B highlights the KEGG pathway enrichment analysis of differentially expressed genes in CaNrf2-TG hearts. Several pathways were significantly enriched, including HIF-1 signaling, glycolysis/gluconeogenesis, PI3K-Akt signaling, MAPK signaling, and the PD-L1/PD-1 checkpoint pathway. Additionally, immune-related pathways such as those implicated in inflammatory bowel disease and rheumatoid arthritis were prominently represented (Figure 1D). These findings collectively indicate that chronic Nrf2 activation disrupts metabolic, immune, and stress response pathways in the myocardium. The x-axis in Figure 1D represents the –log10(FDR), with higher values reflecting greater statistical significance.

**Figure 1:**
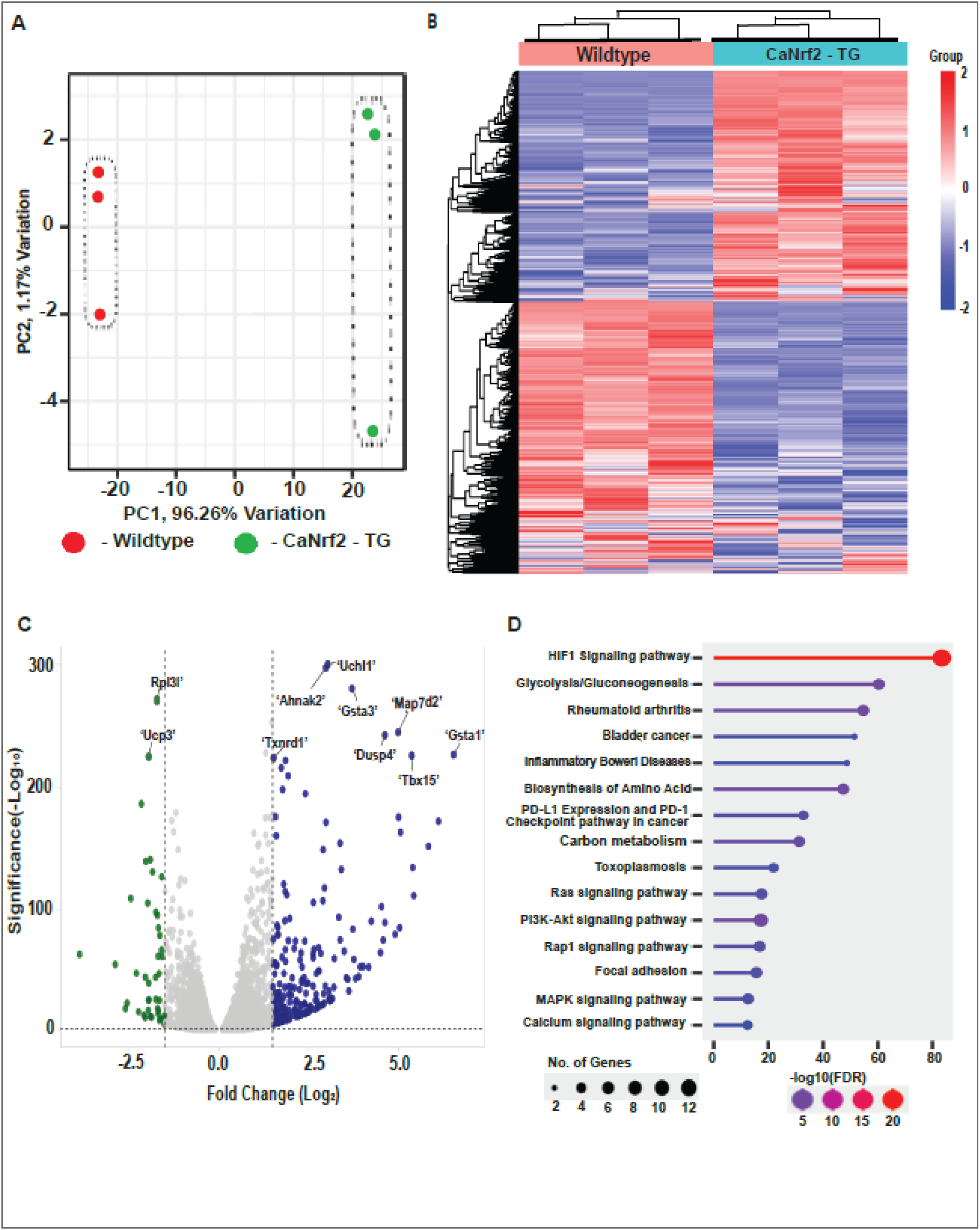
Transcriptomic changes in CaNrf2-TG hearts reveal global Nrf2-driven gene regulation. (A) Principal component analysis (PCA) plot showing strong separation between wild-type and CaNrf2-TG heart samples. (B). KEGG pathway enrichment plot of differentially expressed genes in CaNrf2-TG hearts. Enriched pathways include HIF-1 signaling, glycolysis/gluconeogenesis, PI3K-Akt, MAPK signaling, PD-L1/PD-1 checkpoint, and several immune-related pathways. (C). Volcano plot illustrating the distribution of differentially expressed genes. The x-axis shows log2 fold change and the y-axis shows –log10(p-value). Key upregulated genes include Txnrd1, Gsta1, Ucp3, and Gsta3, while Rpl3l and Ahnak2 are among the downregulated. (D). Summary label aligning data from PCA, volcano, and KEGG enrichment to highlight the coordinated transcriptomic shifts specific to Nrf2 overactivation in the heart.

Figure 1C shows a volcano plot illustrating differentially expressed genes between CaNrf2-TG and wild-type hearts. The x-axis represents log_2_ fold change, while the y-axis represents –log_10_(p-value), highlighting genes with both statistical significance and substantial expression changes. Notably, upregulated genes in CaNrf2-TG hearts include *Ucp1, Txnrd1, Gsta1*, and *Gsta3*, all of which are associated with mitochondrial regulation and antioxidant defense. Conversely, genes such as *Rpl3l* and *Ahnak2* were significantly downregulated. Figure 1D integrates the KEGG pathway enrichment (Figure 1B) and gene-level expression changes (Figure 1C), reinforcing that sustained Nrf2 activation in this model drives widespread alterations in metabolic, redox, and immune regulatory networks.

### 3.2 Mitochondrial Gene Expression Analysis reveals predominant Downregulation of Bioenergetic genes

A focused analysis of genes localized to the mitochondria and its functions confirmed that constitutive activation of Nrf2 in CaNrf2-TG mice results in strong transcriptional reprogramming, as determined by a heat map (Fig 2) with evident segregation between WT and TG samples. Red and green color scales indicating up and down regulated genes respectively explain that the majority of mitochondrial associated genes in TG hearts were suppressed compared to WT. Pathway enrichment analysis confirms that the upregulated genes (Fig) did not show any significant mitochondrial-specific pathways, and terms were restricted to immune and stress responses. This indicates no compensatory activation of mitochondria energy metabolism. Pathway enrichment analysis on downregulated genes (Fig 2) shows significantly enriched pathways associated to central bioenergetics processes such as oxidative phosphorylation, ATP metabolic process, mitochondrial respiratory chain complex assembly and electron transport chain activity. These downregulated genes indicate compromised oxidative phosphorylation function and ATP production.

**Figure 2.**
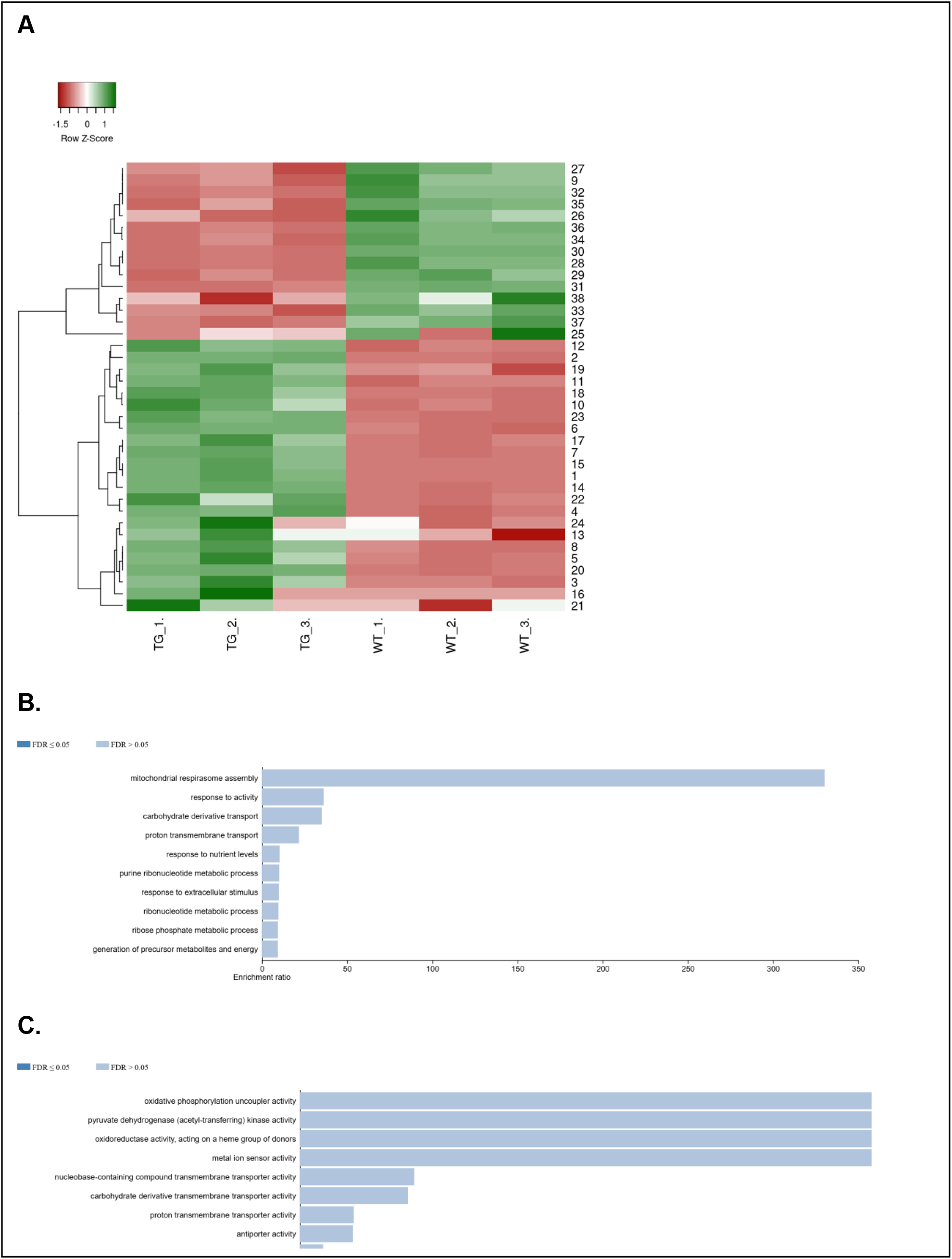
Mitochondrial Gene Expression Analysis Reveals Predominant Downregulation of Bio-energetic Genes in CaNrf2-TG Mice. (A) Heat map showing gene expression profiles in the hearts of wild-type (WT) and CaNrf2-TG (transgenic) mice. The heat map demonstrates a clear segregation between WT and TG samples, with upregulated genes shown in red and downregulated genes in green. The majority of mitochondrial-associated genes in TG hearts are downregulated compared to WT. (B) Bar chart representing pathway enrichment analysis of upregulated genes. The length of each bar indicates the degree of enrichment for each pathway, with blue bars representing pathways with low FDR values (high statistical significance). No significant mitochondrial-specific pathways were identified among the upregulated genes, suggesting that the changes are more related to immune and stress responses rather than mitochondrial energy metabolism. (C) Bar chart representing pathway enrichment analysis of downregulated genes. The length of each bar represents the degree of enrichment, with longer bars indicating more significantly enriched pathways. Significant pathways associated with central bio-energetic processes, such as oxidative phosphorylation, ATP metabolic processes, mitochondrial respiratory chain complex assembly, and electron transport chain activity, are enriched among the downregulated genes.

These findings were consistent with the previous reports (6), Shanmugam et al., 2020 (16), indicates hyperactivation of Nrf2 exerts a predominantly suppressive effect on mitochondrial bioenergetics which in turn explains energetic insufficiency observed in later stages of reductive stress-induced cardiac remodeling.

### 3.3 Functional enrichment analysis reveals distinct biological processes regulated by Nrf2 activation in TG heart at 3 months of age

Differentially expressed genes in CaNrf2-TG and wild-type hearts were analyzed and are presented in Figure 3. Gene set enrichment analysis was performed separately for upregulated and downregulated gene sets, with enrichment significance shown as – log_10_ (adjusted p-value) for each term. For the upregulated gene set, 15 significantly enriched terms were identified across multiple databases, including Gene Ontology: Molecular Function (GO:MF), Biological Process (GO:BP), Cellular Component (GO:CC), KEGG, Reactome (REAC), and Wiki Pathways (WP). Within GO:MF, the top enriched terms included antioxidant activity (Padj = 1.94 × 10^−7^), glutathione transferase activity (Padj = 7.20 × 10^−7^), and oxidoreductase activity (Padj = 1.23 × 10^−7^). GO:BP enrichment highlighted pathways such as cellular detoxification (Padj = 3.34 × 10^−9^), cellular response to xenobiotic stimulus, glutathione metabolism, and response to oxidative stress and lipid hydroperoxides. GO:CC terms were primarily associated with the cytoplasm, extracellular space, and somatodendritic compartments (Fig.3).

**Figure 3:**
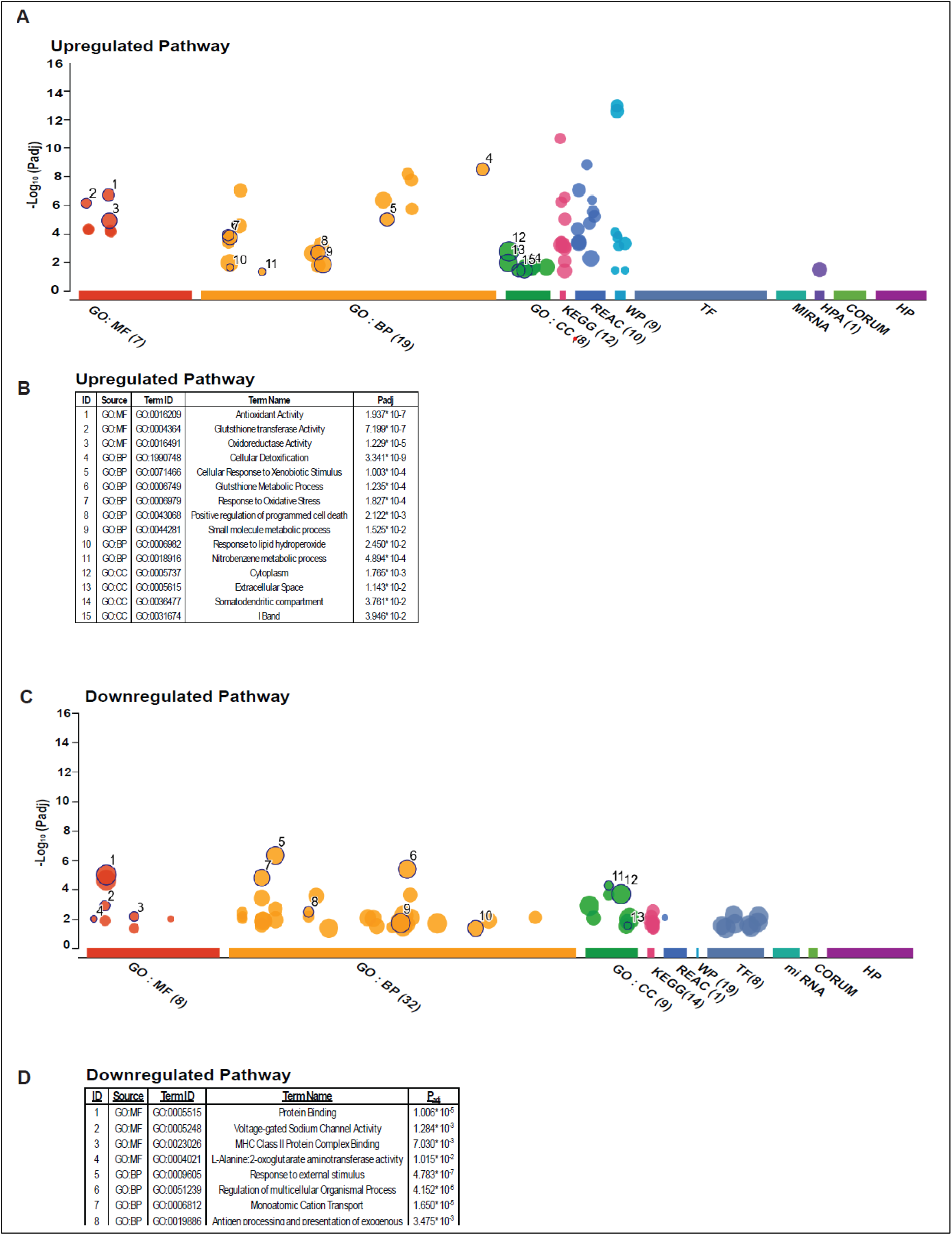
Functional enrichment analysis of upregulated and downregulated genes in CaNrf2-TG and wild-type hearts. Enrichment of upregulated genes across GO Molecular Function (MF) terms. Top enriched terms include antioxidant activity, glutathione transferase activity, and oxidoreductase activity, consistent with known Nrf2 targets. GO Biological Process (BP) enrichment for upregulated genes highlights cellular detoxification, response to xenobiotic stimulus, glutathione metabolism, and oxidative stress response. GO Cellular Component (CC) terms for upregulated genes include the cytoplasm, extracellular space, and somato-dendritic compartments. Enrichment of downregulated genes in GO:MF includes protein binding, voltage-gated sodium channel activity, and MHC class II protein complex binding. GO:BP terms for downregulated genes include responses to external stimuli, antigen processing, and ion transport. GO:CC terms indicate downregulation of components such as the MHC class II protein complex, HCN channel complex, and cell periphery localization.

In contrast, analysis of downregulated genes revealed 13 significantly enriched terms. GO:MF terms included protein binding (Padj = 1.01 × 10^−5^), voltage-gated sodium channel activity, and MHC class II protein complex binding. GO:BP terms involved response to external stimulus (Padj = 4.78 × 10^−7^), regulation of multicellular organismal processes, monovalent cation transport, and antigen processing and presentation. GO:CC terms were enriched for MHC class II protein complex (Padj = 5.47 × 10^−5^), cell periphery, and HCN channel complex. These findings underscore the dual nature of Nrf2 activation, with upregulation of detoxification and antioxidant pathways paralleled by suppression of immune signaling, ion transport, and structural components critical for cardiac function (Fig.3).

### 3.4 Early alterations in gene expression highlight the hypertrophic cardiomyopathy pathway in the TG hearts

Differential gene expressions mapped onto the KEGG Hypertrophic Cardiomyopathy (HCM) pathway (Fig 4) revealed distinct patterns of pathophysiological dysregulation. Several genes involved in extracellular matrix (ECM) interaction and sarcolemma integrity—including ITGA, ITGB, SGCB, SGCD, and DMD—were significantly downregulated. This suggests compromised coupling between the ECM and cytoskeleton, potentially impairing signal transduction and weakening membrane integrity, thereby increasing cardiomyocyte vulnerability to stress and injury.

**Figure 4:**
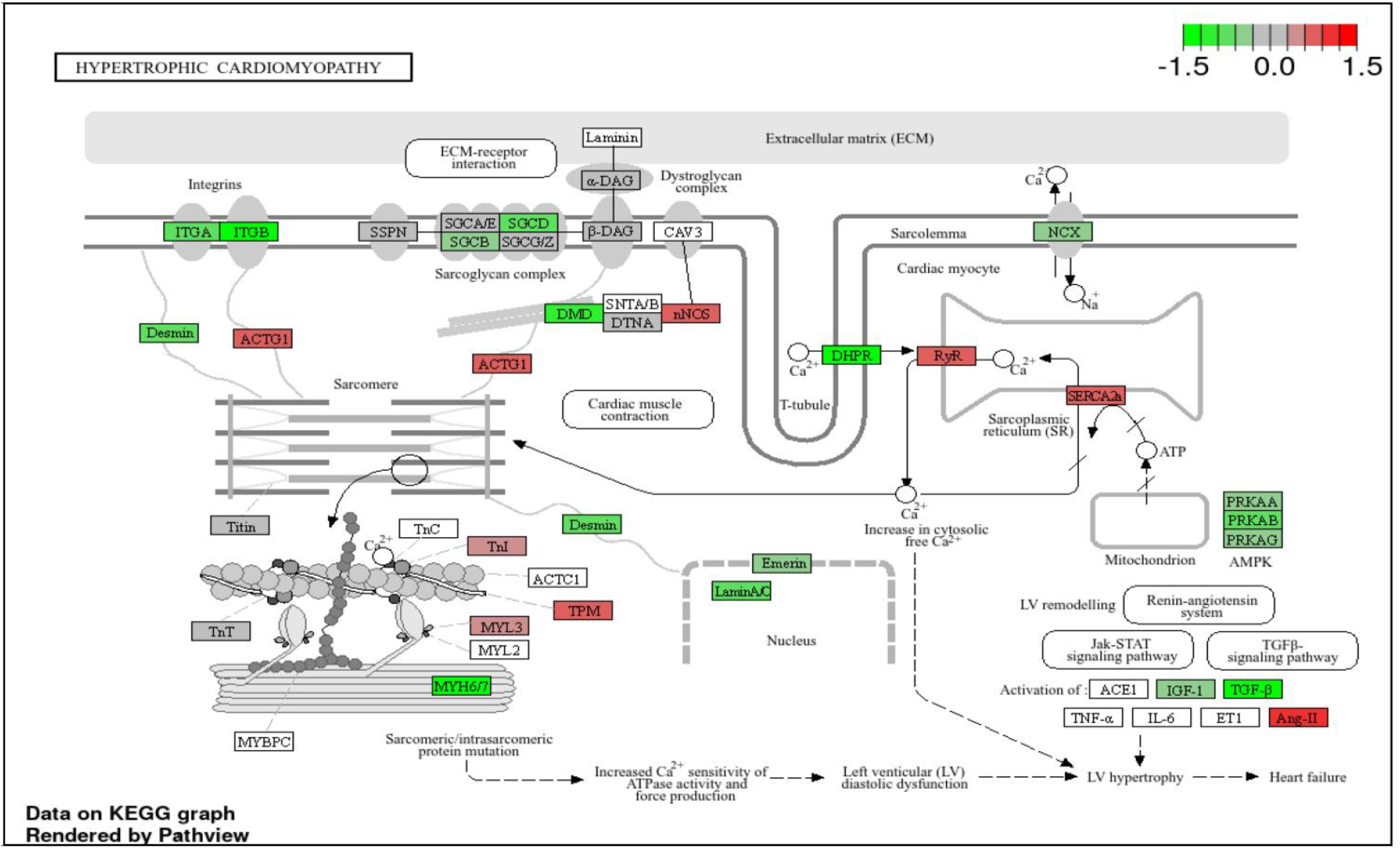
Dysregulation of the cardiac hypertrophic signaling pathway in CaNrf2-TG hearts. Downregulation of extracellular matrix (ECM) and membrane-associated proteins such as ITGA, ITGB, SGCB, SGCD, and DMD. Upregulation of sarcomeric genes (ACTC1, MYL2, MYL3, TPM, TNNI). Calcium signaling alterations include upregulation of RyR and SERCA2a and downregulation of DHPR and NCX. Downregulation of nuclear envelope proteins Emerin and Lamin A/C. Reduced expression of AMPK subunits (PRKAA, PRKAB, PRKAG). Upregulation of pro-hypertrophic and inflammatory mediators (TNF-α, IL-6, ET-1, Ang-II) and downregulation of protective factors (IGF-1, TGF-β).

Within the sarcomere, upregulation of ACTC1, MYL2, MYL3, TPM, and TNNI— encoding key contractile proteins—may reflect a compensatory mechanism to preserve cardiac output. However, concurrent downregulation of MYH6, MYH7, and DES (Desmin), which are essential for force transmission and structural integrity, indicates potential sarcomeric disarray and impaired mechanical stability. Calcium signaling components also exhibited dysregulation. Genes such as RyR and SERCA2a were upregulated, enhancing sarcoplasmic reticulum (SR) calcium release and reuptake, while DHPR and NCX were downregulated, indicating altered calcium influx and extrusion at the sarcolemma. These changes are consistent with elevated cytosolic calcium and increased myofilament calcium sensitivity, contributing to diastolic dysfunction characteristic of HCM (Fig.4).

Further, nuclear envelope proteins Emerin and Lamin A/C were downregulated, suggesting disrupted nuclear-cytoskeletal signaling that may impair transcriptional responses during pathological remodeling. Downregulation of AMPK pathway components (PRKAA, PRKAB, PRKAG) indicates impaired energy sensing and metabolic stress, commonly associated with hypertrophic myocardium (Fig.4). Lastly, altered expression of key signaling mediators was observed. Pro-hypertrophic and pro-inflammatory factors—including TNF-α, IL-6, ET-1, and Ang-II—were upregulated, while protective modulators such as IGF-1 and TGF-β were downregulated. Together, these expression changes (Fig 4) reflect the activation of maladaptive signaling cascades that drive left ventricular remodeling and progression to heart failure in the TG mice as early as 3 months.

### 3.5 Dysregulation of the HIF-1 signaling pathway reveals impaired metabolic adaptation to hypoxia (reductive stress)

Mapping the transcriptomic data onto the KEGG HIF-1 signaling pathway (Figure 4) revealed significant alterations in genes regulating cellular adaptation to hypoxia and metabolic stress. Several genes involved in angiogenesis, the metabolic shift from aerobic to anaerobic pathways, and apoptosis regulation exhibited differential expression, indicating activation of hypoxic signaling. Key upregulated genes included VEGF, ANGPT2, EGF, PKC, PHD, and Camk, reflecting an enhanced hypoxic response that promotes angiogenesis, calcium-mediated stress signaling, and transcriptional adaptation. The induction of HIF-1α target genes such as VEGF and ANGPT2 supports increased angiogenesis under low oxygen conditions. Upregulation of PKC and Camk further suggests activation of calcium signaling pathways and downstream transcriptional events linked to pathological remodeling or survival mechanisms in hypoxic tissue (Fig.5).

**Figure 5:**
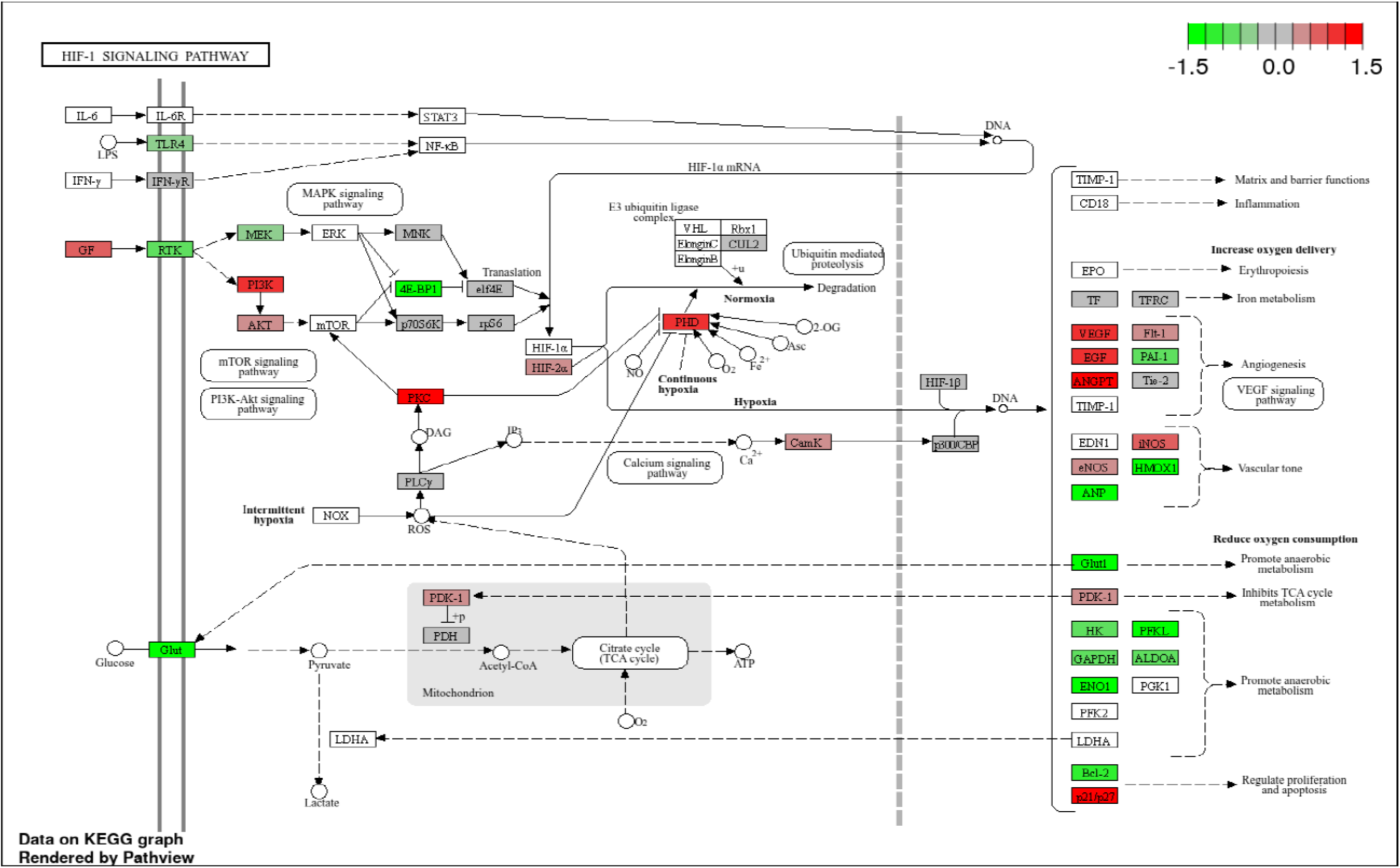
Dysregulation of HIF-1 signaling reveals impaired metabolic adaptation and hypoxia response in CaNrf2-TG hearts. Upregulation of angiogenesis-related genes such as VEGF and ANGPT2, which are canonical HIF-1α targets. Increased expression of PKC, PHD, and Camk. Downregulation of key glycolytic enzymes (Glut1, ENO1, PGKL, ALDOA, GAPDH, HK, LDHA). Suppressed expression of RTK and 4E-BP1, regulators of growth factor signaling and mRNA translation. Expression changes in cell survival and proliferation regulators: upregulation of BCL-2 (anti-apoptotic), and downregulation of p21 and p27 (cell cycle inhibitors). Increased expression of PI3K, AKT, and mTOR and 4E-BP1 downregulation. Upregulation of hypoxia and inflammation-related genes such as EDN1, iNOS, and HMOX1.

Conversely, several genes typically associated with the HIF-1α-mediated glycolytic shift, such as RTK, 4E-BP1, and glycolytic enzymes including Glut1, ENO1, PGK1, ALDOA, GAPDH, HK, and LDHA—were downregulated, indicating an unexpected suppression of metabolic adaptation to hypoxia. Downregulation of Glut1, the primary glucose transporter, suggests impaired glucose uptake and utilization, compromising hypoxia tolerance. Genes involved in apoptosis and cell survival also showed notable changes. Upregulation of the anti-apoptotic gene BCL-2 implies activation of survival pathways in response to hypoxic stress, while downregulation of cell cycle inhibitors p21 and p27 may indicate enhanced proliferation and resistance to apoptosis (Fig.5).

Additionally, Figure 4 highlights upregulation of PI3K, AKT, and mTOR, signaling activation of growth and survival pathways. In contrast, 4E-BP1, a key regulator of translation, was downregulated, suggesting altered translational control and possible feedback inhibition under hypoxic or stress conditions. Pro-inflammatory and vascular tone genes including EDN1, iNOS, and HMOX1 were induced, consistent with pro-hypoxic and vasoregulatory responses aimed at vascular remodeling and nitric oxide-mediated signaling. Collectively, these expression patterns indicate a dysregulated hypoxic response characterized by activated angiogenesis and stress signaling but inhibited metabolic and glycolytic adaptations. This suggests a dissociation between oxygen-sensing pathways and metabolic compensation, potentially contributing to cellular dysfunction during chronic RS caused by constitutively active Nrf2.

## Discussion

This study validates the early and profound impact of Nrf2 activation on the cardiac transcriptome of CaNrf2-TG mice. At 12 weeks of age, CaNrf2-TG hearts exhibit a distinct gene expression profile compared to wild-type controls, as clearly demonstrated by the PCA plot where PC1 accounts for over 96% of total variance. This pronounced separation reflects Nrf2-driven remodeling of cardiac gene expression. Nrf2 is well-known as a master regulator of cellular defense, especially through the induction of antioxidant and detoxification genes (12). However, chronic overactivation of Nrf2, as modeled in CaNrf2-TG mice, can shift the redox balance towards a hyper-reductive state—referred to as reductive stress (RS)—which disrupts cellular homeostasis and promotes pathogenic effects, particularly in energy-demanding organs like the heart (6-9, 13, 14).

Understanding disease progression requires considering the temporal transcriptomic changes in CaNrf2-TG hearts. We recently reported that at 4 weeks, early molecular alterations emerge, including upregulation of canonical cardiac stress markers (Nppa, Nppb) and heat shock proteins (Hspa1l, Dnajb40), likely reflecting early compensatory or cytoprotective responses (13). Simultaneously, downregulation of genes critical for endoplasmic reticulum (ER) homeostasis and protein folding (Kfbp4, Ppil1, Cav1) suggests early strain on cellular stress adaptation mechanisms (13).

By 12 weeks, these disruptions become more pronounced. Key ER chaperones such as Hspa5 (BiP/GRP78), Hspd1, and Hsph1, essential for ER integrity and protein folding, are downregulated, alongside other ER proteostasis components like Calr, Pdia3, and Canx. This pattern indicates a failure of adaptive stress responses and progression toward maladaptive remodeling (13, 15). Interestingly, the transcript levels of Pdi, an ER enzyme, is upregulated at this stage, possibly representing a cellular effort to restore ER function.

By 24 weeks, the disease phenotype is fully established (16, 17). Significant changes occur in approximately 35 ER-related genes. Notably, extracellular chaperone Clu (Clusterin) is downregulated, while Vcp, involved in ER-associated degradation (ERAD), is upregulated, highlighting a shift from repair to degradation pathways and chronic stress (16). Throughout this timeline, antioxidant genes regulated by Nrf2—such as Nqo1, Gclc, and Gstm—remain persistently overexpressed, indicating the heart is not under oxidative stress from insufficient antioxidants but rather suffers from a reductive-redox environment (16). While antioxidant overexpression is initially protective, sustained elevation under unstressed conditions can disrupt normal redox signaling and proteostasis, contributing to an impaired PQC-related pathology.

These findings align with pharmacological models of chronic Nrf2 hyperactivation. High-dose, prolonged treatment with Nrf2 activators like sulforaphane or N-acetylcysteine mediated glutathione (GSH) augmentation induces similar maladaptive gene expression changes, including induction of stress markers and suppression of ER protein folding genes, followed by upregulation of ERAD and protein degradation pathways alongside decreased critical chaperones (18, 19). In contrast, lower or intermittent dosing promotes adaptive antioxidant responses without triggering reductive stress.

KEGG and GO enrichment analyses further support these conclusions. CaNrf2-TG hearts show marked alterations in immune response pathways (PD-L1/PD-1, MHC-II antigen presentation), cellular signaling (PI3K-Akt, MAPK, HIF-1), and metabolism (glycolysis/gluconeogenesis) (4, 5, 20). Downregulated genes reveal broad suppression of protein folding, immune regulation, and cellular communication needed for homeostasis, whereas upregulated genes enhance antioxidant and detoxification functions (3, 13, 21). While hyperoxia and elevated ROS/RNS levels cause oxidative stress, a reduction of ROS to sub-physiological levels leads to reductive stress, mimicking a hypoxic state. In this study, we found that under chronic hypoxic conditions (i.e., reductive stress), HIF1α signaling is insufficiently activated, potentially triggering pathological cardiac remodeling. We propose a novel hypothesis that reductive stress– induced impairment of HIF1α signaling may represent an unresolved mechanism underlying proteotoxic cardiac remodeling.

## Conclusions

Constitutive Nrf2 activation in CaNrf2-TG hearts induces early and sustained transcriptomic shifts, converting protective antioxidant signaling into maladaptive reductive stress. This disrupts ER proteostasis, cellular signaling, and metabolic regulation, driving progressive cardiac pathology. The 12-week stage marks a crucial transition point where compensatory mechanisms collapse, highlighting a therapeutic window to counteract Nrf2-induced remodeling. These findings emphasize the context-dependent nature of Nrf2 activity and the need for precise modulation to prevent reductive stress–driven cardiomyopathy.

## 2. Materials and Methods

### 2.1 Animal model and Study design

Cardiac-specific, constitutively active Nrf2 (CaNrf2-TG) transgenic mice were utilized as described in our recent study (17). Littermate wild-type (WT) mice were utilized as controls. Male mice were analyzed at age of 12 weeks, based on prior findings it has been identified that this time point as a transitional phase in disease development (15). All experimentation involving experimental animals was approved by the Institutional Animal Care and Use Committee at the University of Albama at Birmingham and conducted according to NIH guidelines.

### 2.2 RNA isolation and sequencing

RNA was isolated from 12 weeks old male WT and CaNrf2-TG mouse hearts using RNeasy Mini kit (brand and catalogue) according to manufactures protocol. RNA isolation and sequencing was performed as described in our previous study (22). RNA quality was confirmed using Nanodrop spectrophotometer and bioanalyzer. Poly (A) enriched RNA was used to prepare libraries using the Illumina Stranded mRNA Prep and indexed libraries were pooled and sequenced on Illumina NovaSeq.

### 2.3 RNA sequence Alignment and Differential Expression Analysis

The data generated was subjected to further analysis by removing low quality bases and adaptor contamination using tools like FastQC and Trimmomatic. The filtered data was then mapped to Mus musculus genome from the ensembl database. DESeq2 tools were used for further analysis to obtain the differentially expressed genes and the fold change information. Differentially expressed genes (DEGs) were then further filtered out by keeping the (log2 fold change) >1 and p-value < 0.05 as the cutoff criteria. Functional enrichment of the curated genes were then performed using a combination of tools and databases like KEGG, GO, R eactome and WikiPathways. Volcano plots and enrichment bar plots were generated using R packages ggplot2 and clusterprofiler.

### 2.4 Principal Component Analysis (PCA) and visualization

PCA was conducted using variance stabilized transformed counts obtained from DESeq2. The PCA plot was generated using the ggplot2 R package to visualize sample clustering and separation between WT and CaNrf2-TG groups.

### 2.5 Hypertrophic Cardiomyopathy and Mapping of HIF-1 Pathway

To identify the functional significance of transcriptomic changes, DEGs were overlapped onto the KEGG hypertrophic cardiomyopathy (HCM) and HIF-1 signaling pathways. Genes participating in sarcomere function, calcium handling, extracellular matrix (ECM) integrity, metabolic regulation and hypoxia response were checked for expression variations. Mappings were created with the KEGG mapper tool.

## 2.6 Data Availability

**The Data generated from the NExt Generation Sequencing on Illumina NovaSeq will be made available in NCBI**.

## Supplementary figures legends

**Figure 1:** Quality assessment and reproducibility of RNA-seq data from wild-type and CaNrf2-TG hearts. (A) Classification of detected transcripts into coding genes, pseudogenes, and long noncoding RNAs. (B) Boxplots of transformed expression values showing comparable distributions between wild-type (WT) and CaNrf2-TG samples. (C) Density plots of transformed expression values confirming effective normalization across biological replicates. (D) Total raw read counts (millions) for each replicate, demonstrating consistent sequencing depth across WT and CaNrf2-TG hearts. (E) Scatter plot comparing transcript expression between WT and CaNrf2-TG groups (R = 0.92, p < 0.01). (F) Scatter plot showing near-identical expression profiles between two independent CaNrf2-TG replicates (R = 1.0, p < 0.01). (G) Scatter plot comparing two WT replicates (R = 1.0, p < 0.01), confirming high reproducibility of RNA-seq data across samples.

**Figure 2:** Protein-Protein interaction network of cardiac hypertrophy highlighting differentially expressed genes in CaNrf2-TG hearts. (A) Genes within the hypertrophy signaling pathway are color-coded by differential expression: red indicates upregulation and green indicates downregulation in CANrf2-TG hearts. (B) The diagram illustrates the organization of key molecules involved in stress signaling, transcriptional regulation and structural remodeling during cardiac hypertrophy with gene-level changes overlaid to show Nrf2 activation affects specific nodes in the pathway.

**Figure 3:** Pathway enrichment analysis of differentially expressed genes in CaNrf2-TG versus wild-type hearts. Pathway enrichment analysis of significantly altered transcripts (adjusted p < 0.05, |log_2_FC| ≥ threshold) identified multiple Nrf2-associated signaling cascades. (A) Upregulated pathways include glutathione metabolism, drug metabolism– cytochrome P450, metabolism of xenobiotics, and other antioxidant defense pathways, consistent with enhanced redox buffering and detoxification capacity driven by Nrf2 activation. (B) Downregulated pathways are enriched for mitochondrial oxidative phosphorylation, fatty acid metabolism, and protein processing in the endoplasmic reticulum, indicating suppression of metabolic and proteostatic networks. Together, the data suggest that cardiac-specific Nrf2 overexpression leads to global transcriptional reprogramming characterized by activation of cytoprotective and redox-regulating processes while concurrently repressing energy metabolism and proteostasis-related pathways.

