## Supplementary figures and images for "Early Transcriptomic Signatures of Reductive Stress Cardiomyopathy Reveal Insufficient HIF-1α Signaling"

### Supplemental figures

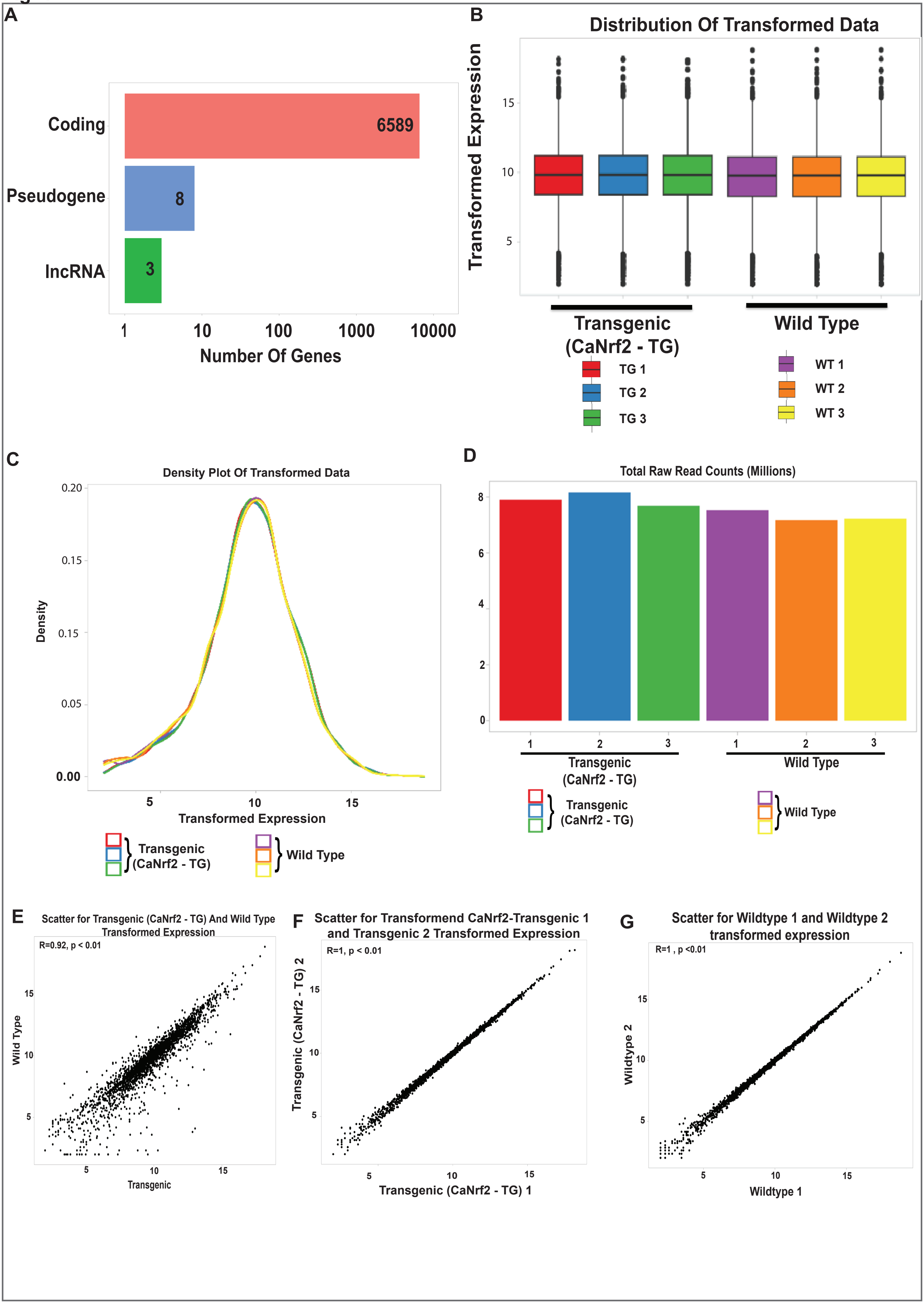

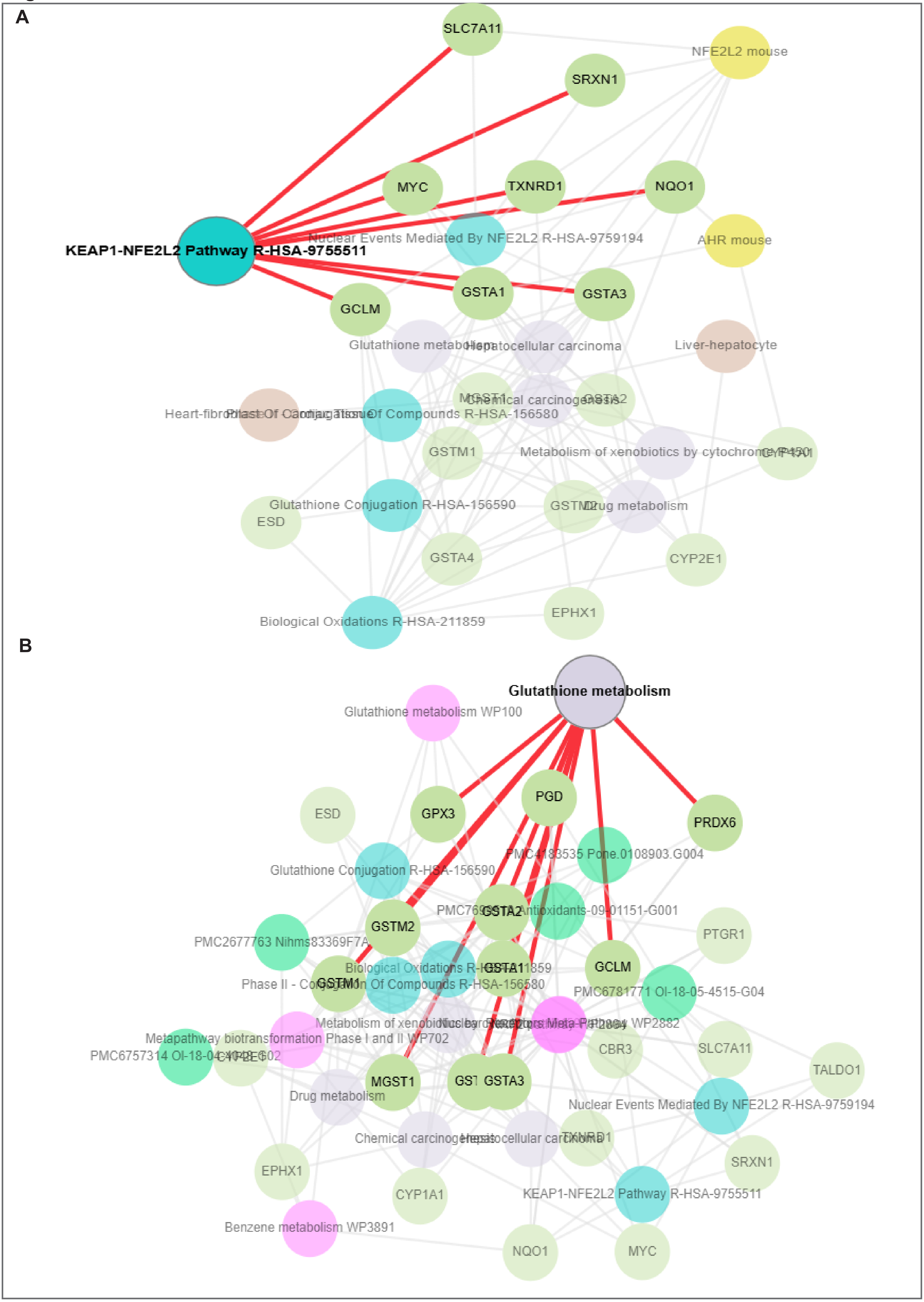

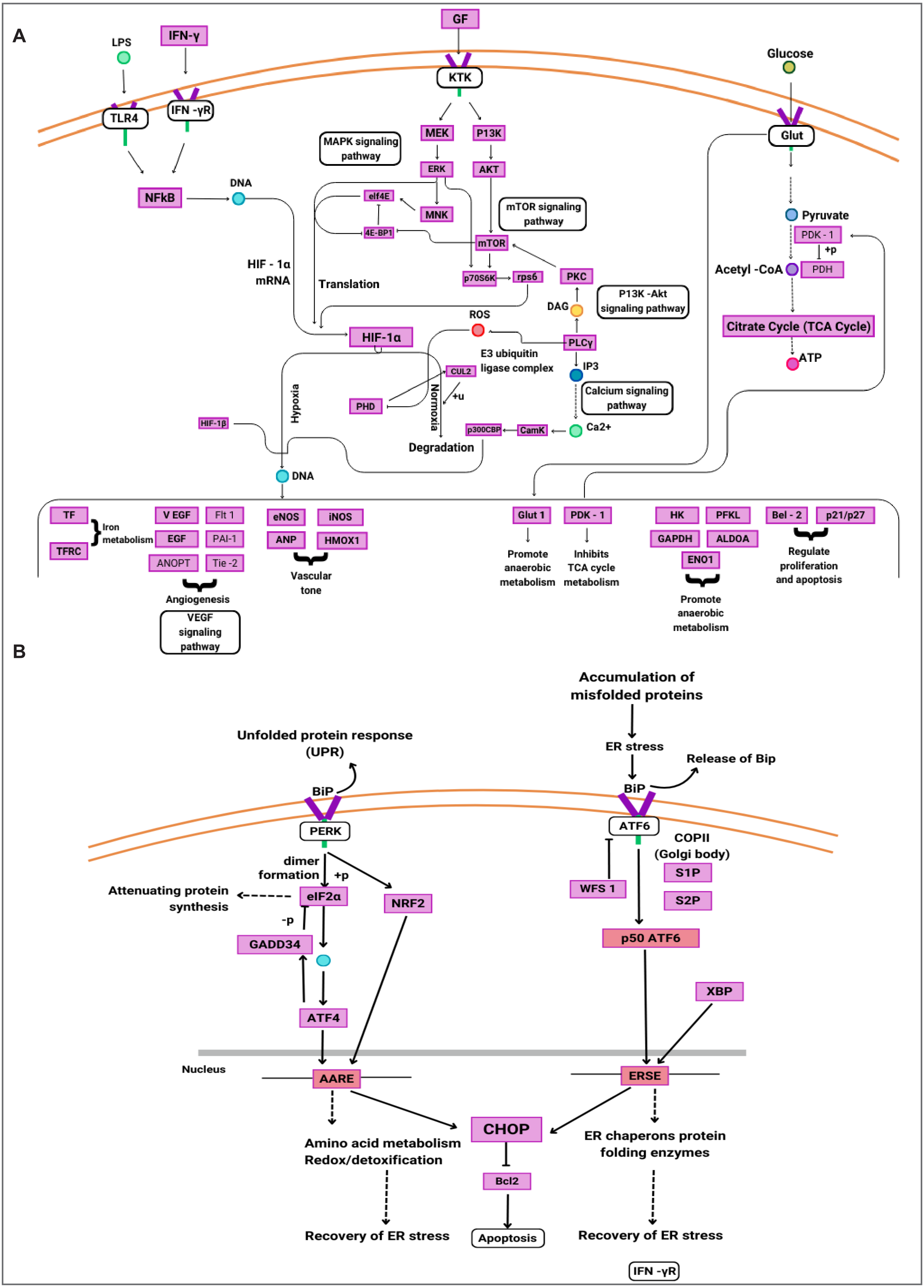
